# Parsing of compositions and microstructure characteristics for rust-spots of pear pericarp

**DOI:** 10.1101/142877

**Authors:** Lijun Nan, Shaobo Chen, Yashan Li, Ya Liu, Ying Jiang, Yan Yang, Chengdong Xu, Guogang Chen

## Abstract

The commercial value of Kurles pears pericarp, a popular and favored fruit for its unique aroma and refreshingly crisp texture, had sharply decreased because of a rust breakout of the beloved pear in China during the atmosphere-controlled storage. High performance liquid chromatography (HPLC) and liquid chromatography-mass spectrometry (LC-MS) were used to analyze rust spots on the pericarp of Kurle pears. Therefore, the chemical compounds of four various eluates, originating from the rust-colored substance collected from the pears pericarp effected, were identified successfully for the first time, which were just rhein, aloe-emodin, chrysophanol and emodin, respectively. Taken together with microstructure characteristics for rust-spots of Kurle pear pericarp, it was no doubt that these eluates were the main factors affecting the rust spots on the pericarp of the Kurle pears during the atmosphere-controlled storage, which was a sign and consequence resisting the undesirable stress of the external environment.

## Introduction

Besides of the grapes and apples, the pears are also the most popular temperate fruit species in China at present. Whereas there was an irrefragable fact that the pear industry had become financially burdensome recently due to be increasingly ruined by a number of the fungi diseases. In Europe, there was many reports about the pears having been perplexed by some sickness, such as the scab, powdery mildew, brown spots, fire blight, and other fungal diseases. Nevertheless, the pears had been suffering predominantly from the scab and rust caused by the black spot disease in Asia (Itai *et al.,* 2012), which had become the most significant trouble in term of the future of pear industry needing confront with the bravery.

As a rare species in the world, the Kurle pears were desired for the pure fragrance, refreshing juice, and tender quality and so on. The Kurle pears were frequently exported and highly sought after all throughout Southeast Asia and Europe nowadays. However, the frequent appearance of the rust spots on the pear cuticles had recently depreciated the commercial value of the fruits in a substantial way. Few studies had examined in detail this recent phenomenon affecting the exterior quality of this prized fruit (Liu *et al.,* 2013; Stanislaw *et al.,* 2015).

There were many methods available for the material analysis and identification until now, such as the chemical reaction, liquid chromatography (LC), gas chromatography (GC), thin layer chromatography (TLC), visible spectroscopy (UVC), liquid chromatography mass spectrometry (LC-MS), and nuclear magnetic resonance spectroscopy (NMRS). It was essential for us to choose an appropriate method analyzing and identifying the properties of the matter tested. To obtain the most accurate information about the experimental material, two or more methods of the analysis and identification should usually be adopted simultaneously. Thus, LC-MS and LC, as the combined methods, were chosen meanwhile in this trial to analyze and identify the rust spots of the Kurle pear, which would provide the theoretical basis as for the relevant technology controlling the rust spots on the Kurle pear pericarp.

In term of every creature, the cell was the basic unit of the organ and tissue performing the physiological function. Under the atrocious environmental stimulation inevitably, the combined action between the broken cell wall and the lignification reaction inside the cell formed the cork cambium from the pear peel of the rusty spot (Clairmont *et al.,* 1992). The thin epidermal cells of the fruit peel were damaged easily and developed quickly the cork layer under the stimulation of the severe environment, while the cork cell accumulating on the skin developed the terrible rusty spot. As we all known, there were a lot of the rough endoplasmic reticulum in the plant epidermal cell, which was the source of many organelles, such as the vacuoles and golgi apparatus, whereas the material transportation between the cells could been regulated constantly and steadily by the biochemical process of the endoplasmic reticulum. Two experiments of Teasdale and Jackson (1996) and Xie *et al.* (2009) had demonstrated that the endoplasmic reticulum was not obvious degradation role during the fruit maturation excluding the final stages of the overripe. In view of these conclusions, the analysis of the ultramicroscopic structure, as far as the rusty spot area and intact cells of the korla fragrant, could lay a foundation for the identification of the rusty spot material.

## Materials and methods

### Sampling and treatment of fruits

The Kurle pears used in this study, kept in atmosphere-controlled storage, were sampled according to the concrete experiment demands on Novermber 22, 2015 from Kurle city, China, respectively.

The fresh - keeping storehouse with the atmosphere - controlled function, 50 m^2^, stored 10,000 Kg of the Kurle pears, which were kept in 4 m hight, 50 cm altitude per storey of goods shelf, and a layer of pear in every storey of good shelf, of storage rack. 10 Kg parent population of fruits per replication were sampled randomly from the middle and four sides of the storeroom. Classification, 4 sorts, was carried out according to the shape and color of the rust spot, excluding a sigle of the same category without the defect of the fruit. 10 fruits, the similar shape and color of the rust spot, were pooled to obtain one sample for the analysis. All factors was repeated three times.

The 5 g samples of the rust spots per sort above, treated with a 125 mL ethanol (70%) solution after weighed accurately, were extracted from the surface of the pears. Then, the liquid extracted was concentrated immediately to a ratio of 1: 4 after being eluted with the petroleum ether - acetone and petroleum ether - ethyl acetate gradient elution further, which were named elution 1, elution 2, elution 3 and elution 4, respectively. Additionally, the reagents of the chloroform and petroleum ether were added as another extraction liquid (EL) of the extraction agents with water (EL : water =1: 1).

Finally, the rust spots liquids of the pears above were separated and purified with the method of the column chromatography, respectively. The procedure was listed as follows. Firstly, the petroleum ether extract was washed by the mixtures of the petroleum ether and acetone with two different ratios of the eluates (4: 1, 5: 1), respectively. The elution 1 was collected in the cyclone separator of the extraction unit. Secondly, the chloroform extract was washed with the mixtures of the petroleum ether and ethyl acetate with three different gradients of the eluates (5: 1, 4: 1 and 2: 1), respectively. In addition, the samples, namely elution 2, 3 and 4, were also collected in the cyclone separator of the extraction units, respectively. It was vigilant that the liquid level, not less than the upper surface of the quartz sand, was always ensured during the chromatographic separation experiment, and the cock of the chromatography column should be closed in time before the samples were added. In addition, it should be noticed that the dropper should not bring into contact with the upper surface of the quartz sand until the samples were injected continually, while the cock of the chromatography column must be immediately opened after finishing the sample injection. At alst, the eluting agent was collected smoothly and continuously within a 2 - mL bottle numbered at the rate of 5 - 8 drops per minute, respectively, and prepared for the test of the purity.

### Samples analysis of LC - MS

The samples, including elution 1, elution 2, elution 3 and elution 4, were analyzed with LC - MS, respectively. The analysis conditions were as follows: mobile phase (firstly 0.1% of 100% formic acid, 10 min; followly 0.1% miscible liquids of 40% acetonitrile + 60% formic acid, 20 min; finally 100% acetonitrile, 8 min), column temperature (45 °C), wavelength (200 - 400 nm), flow velocity (0.3 mL/min), sample volume (1 μL); ion form (ESI^-^ and ESI*+*), capillary voltage (3.0 and 3.5 kVolts), cone voltage (20 and 20 Volts), ion source temperature (100 and 100), desolvation temperature (250 °C and 250 °C), removal of solvent gas flow (500 and 500 lit/hr), conical flow (50 and 50 lit/hr), collision energy (15 and 15 Volts), mass range (100 - 1500 and 100 - 1500 m/z), detector voltage (1700 and 1600 Volts).

### Samples analysis of LC

The another samples, elution 1, elution 2, elution 3 and elution 4, were analyzed with LC referring to the standard mixture of both (Li *et al.* 2011) and (Purnhauser *et al.* 2011). The experimental conditions were as follows: mobile phase (Methanol: 0.1% Phosphoric acid) = 85: 15, column temperature 25 °C, wavelength 254 nm, flow velocity 0.3 mL/min, sample volume 20 μL. A qualitative analysis was also performed in accordance with the standard mixture and retention time of the four elution solutions.

### Ultrastructure of fragrant pear cell

The intact parts of the Kurle pears was taken as control to illuminate and ascertain the microstructure characteristics of the rusty spot of the experimental material during the atmosphere - controlled storage. 3% glutaraldehyde solution was configured by 0.05 mol/L phosphate buffer (pH 6.8) first. The peel and pulp of the fragrant pear, cut into 1 mm x 2 mm (longth x width) wafer by the scalpel, was fixed 24 h with 3% glutaraldehyde solution under 4 □ before being rinsed 3 - 4 times with phosphate buffer (pH 6.8). After that, the samples, fixed 2 h by 1% osmic acid solution at room temperature, followed 30%, 50%, 60%, 70%, 80%, 90% and 100% ethanol dehydration in gradient, 15 minutes each time after being also washed 3 - 4 times by the phosphate buffer (pH 6.8), respectively. The materials above were successfully replaced three times with the acetone further, then embedded both about 2 h by the embedding medium (acetone: epoxy resin = 3: 1) and over 4 h with the medium (acetone: epoxy resin= 1: 1), respectively, prior to the polymerization of the temperature - keeping box at 60 ┚ for 24 h. Finally, the polymerized samples were sliced into the slices (about 60 nm of thickness) with the ultra - thin slicing machine in order to the observation and photograph under the transmission electron microscope.

### Instruments and equipments

Instruments and equipments in the trial was readied as follow:

Hitachi H - 600 transmission electron microscope, OLYMPUS CX21FS1 optical microscope, 705902 ultra - thin slicing machine (leica in Australia), HPLC (LC - 2010A, Shimadzu, Japan), LC - MS (LCMS8030, Shimadzu, Japan), chromatograph (Waters acquity uplc), detector (Waters acquity pda), and analytical column (BEH, C18, 2.1×100 mm 1.7 μm).

### Reagents

Instruments and equipments in the trial included: methanol (GR, Fair Lawn, USA), acetonitrile, rhein, chrysophanol, aloe emodin and emodin (GR, Sigma, USA). phloroglucinol, safranine, 95% ethanol, osmic acid, iodine, potassium iodide, glutaraldehyde, epoxy resin and phosphoric acid (AR, BASF, Tianjin), formic acid (AR, Xi’an Chemical Reagent Factory).

## Results

### LC-MS analysis of Elution 1

According to the mass spectrum (Fig. 1), the molecular weight of the elution 1 was 284. The mass spectral information, m/z, was as follows: 283 [M^+^], 239 [M^+^ - COOH], 211 [M^+^ - COOH – CO], 183 [M^+^ - COOH – CO - CO]. In comparison with the prototype HPLC information under the same chromatographic conditions, the identification result showed that the retention times were fairly consistent in regard to the elution 1 (25.055 min) and rhein standard (25.068 min) (Fig. 5a and 5b). Based on the analysis above, it was quite obvious that the elution 1 could be identified as rhein, and molecular structure of which was displayed in Fig. 6a.

**Fig. 1.**
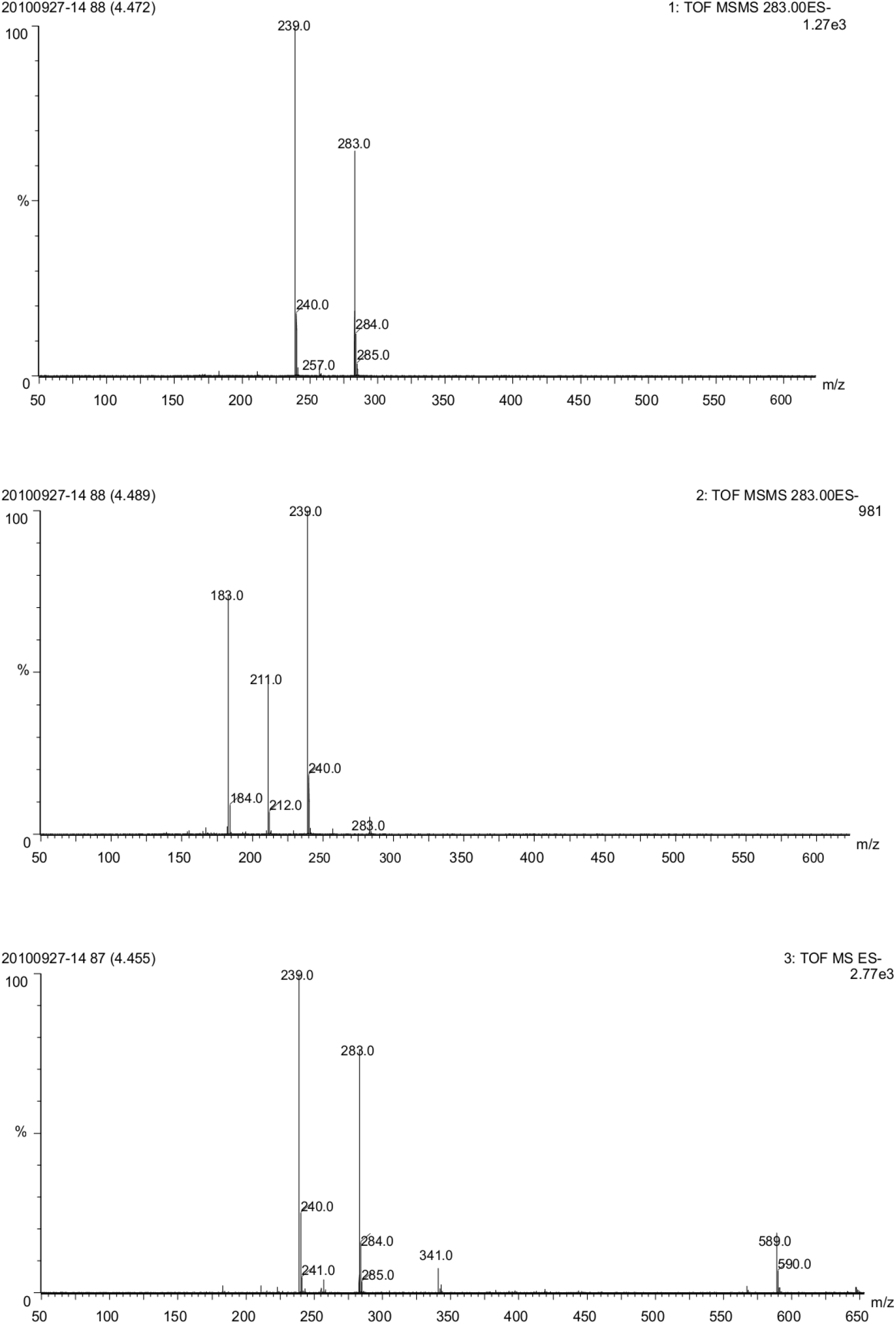
Mass spectrum for elution1 of the rust spots

### LC-MS analysis of Elution 2

The molecular weight of the elution 2, 270, could be also seen in Fig. 2, and the results of the mass spectral information (m/z) included: 270 [M^+^], 269 [M^+^ - H], 241 [M^+^ - CO - H], 223 [M^+^ - 2H - COOH], 195 [M^+^ - 2H – COOH - CO], 183 [M^+^ - COOH – CO - CH_2_]. Through the prototype HPLC information under the same chromatographic conditions, the retention times of the identification result were fairly the same as the elution 2 (18.998 min) and aloe emodin standard (18.981 min) (Fig. 5a and 5c). Based on the fact above, the elution 2 was ascertained as aloe emodin, and molecular structure of which was showed in Fig. 6b.

**Fig. 2.**
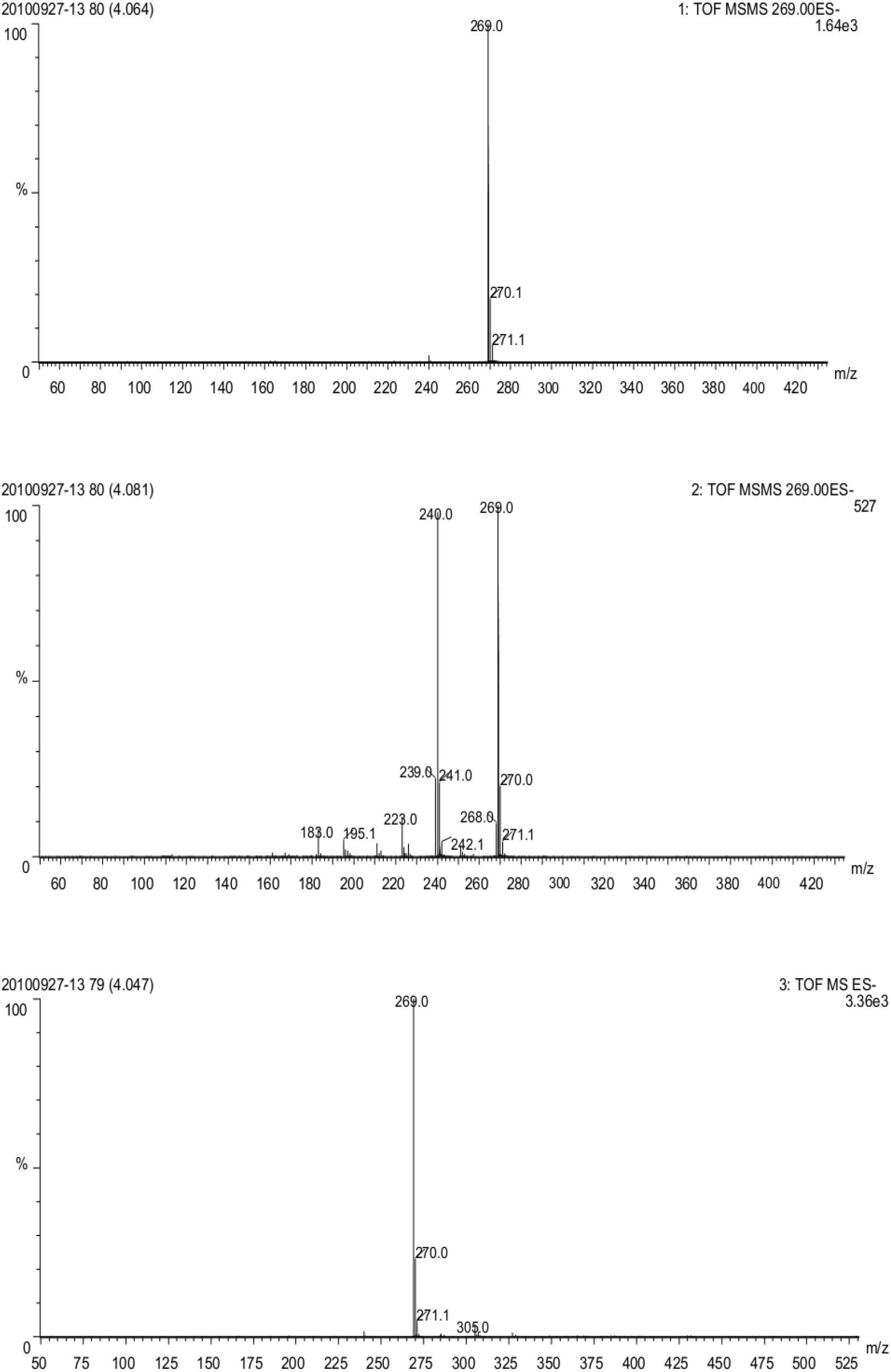
Mass spectrum for elution 2 of the rust spots

### LC-MS analysis of Elution 3

According to Fig. 3, the molecular weight of the elution 3 was 254. The mass spectral information was as follows (m/z): 254 [M^+^], 253 [M^+^ - H], 225 [M^+^ - COH], 162 [M^+^ - 2COH - 2OH], 158 [M^+^ - COH - CO -3CH]. The prototype HPLC information was referred to as the control under the same chromatographic conditions, the fairly consistent retention times, the elution 3 (44.028 min) and chrysophanol standard (43.948 min), were found in Fig. 5a and 5d. Therefore, the elution 3 was affirmed unquestionably as the chrysophanol, and the molecular structure of which was showed in Fig. 6c.

**Fig. 3.**
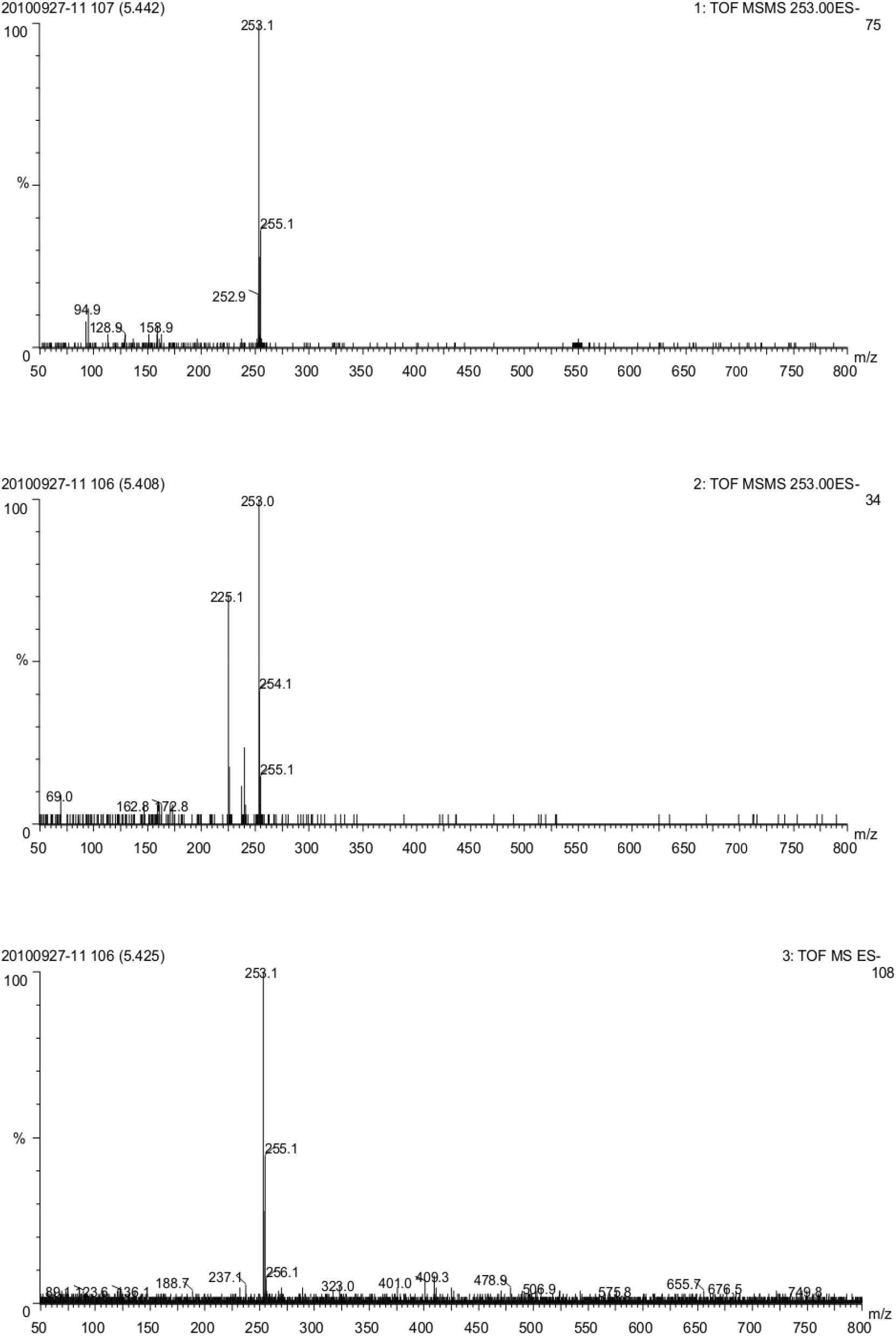
Mass spectrum for elution 3 of the rust spots

### LC-MS analysis of Elution 4

It was satisfied that the molecular weight of the elution 4, 270, could be also obtained based on Fig. 4. The mass spectral information was distinct (m/z): 270 [M^+^], 269 [M^+^ - H], 241 [M^+^ - CO - H], 225 [M^+^ - CO - OH], 197 [M^+^ - 2CO - OH]. Analyzing the prototype HPLC information, the retention times of both the sample (35.177 min) and the standard reagent (35.022 min) under the same chromatographic condition. Through the summary on the above analysis, the elution 4 was identified as the emodin (Fig. 5a and 5e), and the molecular structure of which was displayed in Fig. 6d.

**Fig. 4.**
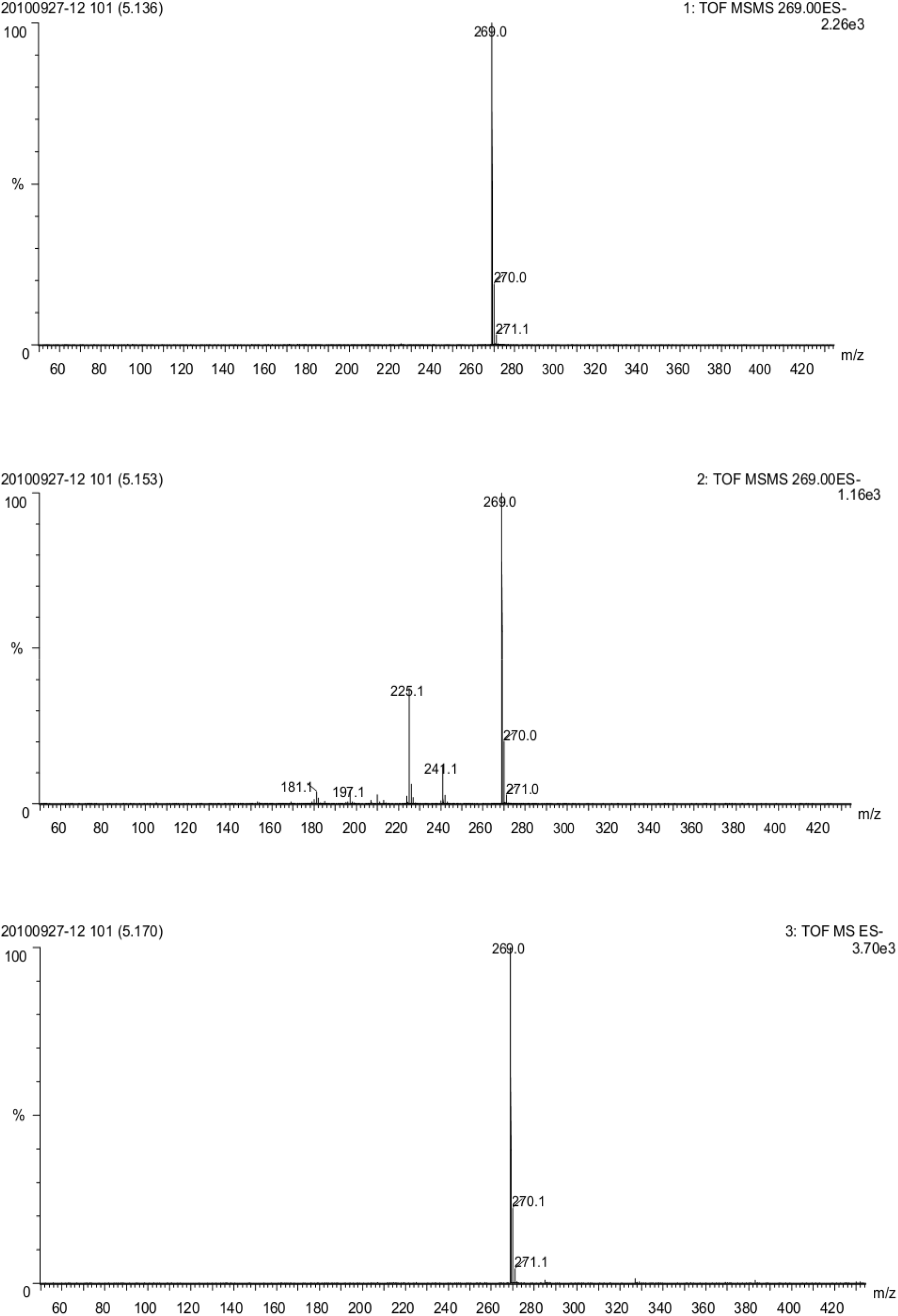
Mass spectrum for elution 4 of the rust spots

## Discussion

### LC - MS analysis of Elution 1

From the more careful check, there was a the most pleasant surprise that the result was in accordance with the previous data about the mass spectrum of the rhein (Doğmuş-Lehtijärvi *et al.,* 2009; Brewer *et al.,* 2009; Doğmuş *et al.,* 2010; Mathias *et al,* 2011).

Rhein was thought of as the anthraquinone compounds (Cben *et al.,* 2011). Anthraquinone ring containing two hydroxyl and a carboxyl possessed the stronger polarity and the characteristics of a strong oxidation reduction. Therefore, the rhein was not stable very much, and decomposed easily by the oxidation - reduction reactions. Rhein, based on the fact above, should be fond of the anaerobic environment. The controlled atmosphere storage method, adopted in this experiment created exactly an anaerobic environment for the fragrant pear, could resist effectively the oxidation of the external environment. As we all known, the rhein could provide some roles, such as the antibacterial and scavenging free radicals (Rodríguez *et al.,* 2007). Therefore, the rhein was advantageous to the storage of the fragrant pear owe to protecting the organization from the damage. Some measures, such as increasing the light intensity, raising the temperature and improving pH, could accelerate the degradation rate of the rhein with the degradation reaction of the first order kinetics process. When the pH > 8, the reaction rate of the rhein degradation would increase rapidly. So the rhein was more suitable for a cool environment avoiding the violent glare and stronger alkaline environment. This fragrant pear with a certain acidity, adopting the controlled atmosphere storage, was conserved in the lower temperature, oxygen free or micro oxygen environment in the trial, therefore, the rhein could be easily detected inevitably, which was also the embodiment of the korla fragrant pear with a strong antioxygenic property.

### LC - MS analysis of Elution 2

Beyond all reasonable doubt that the results of the mass spectral information in Fig. 2 was consistent with the previous datas for aloe emodin (Crampton *et al.,* 2009; Doğmuş‐Lehtijärvi *et al,* 2009; Brewer *et al.,* 2009; Douglas *et al.,* 2010; Mathias *et al,* 2011; Itai *et al.,* 2012).

Aloe emodin belonged to anthraquinone compound, which usually existed in Chinese herbal medicine, such as aloe, rhubarb and cassia seed. This experiment also detected the aloe emodin in the korla fragrant pear. The effective antibacterial ingredients of the aloe emodin possessed some obvious features, such as the anti - tumor activity, antibacterial activity and immunosuppression and purgation (Li *et al.,* 2012; Hu *et al.,* 2014). Therefore, the aloe emodin in the korla fragrant pear had played a part in the certain antibacterial activity to the pear fruit, the body function could be improved obviously after the korla fragrant pear with the aloe emodin was eated. The pharmacological activity of the aloe emodin was related closely to the chemical structure of the aloe emodin itself. An anthraquinone ring and two phenolic hydroxyl of which determined the function of the biological activities including the removal of the oxygen free radicals, antitumor and other aspects (Ruie *et al.,* 2014). Another study demonstrated yet that the aloe emodin could enhance the phagocytosis of the macrophage, and induce the mRNA expression of the cytokines including interleukin in the white blood cells, except for the tumor necrosis factor and interferon (Yu *et al.,* 2006) so as to enhance the body’s immune function. Therefore, the aloe emodin contained could protect the skin of the korla fragrant pear from the erosion of the foreign organisms. Under the condition of the controlled atmosphere storage in this trial, the skin of the korla fragrant pear would generate spontaneously the aloe emodin, which was a reflection of the adaptability characteristic to resist the external environment for the korla fragrant pear.

### LC - MS analysis of Elution 3

A delirious and unavoidable fact was that the mass spectral information in Fig. 3 and the prototype HPLC information in Fig. 5a and 5d in the trial were in accord with the previous data for chrysophanol (Manjunatha *et al.,* 2008).

**Fig. 5.**
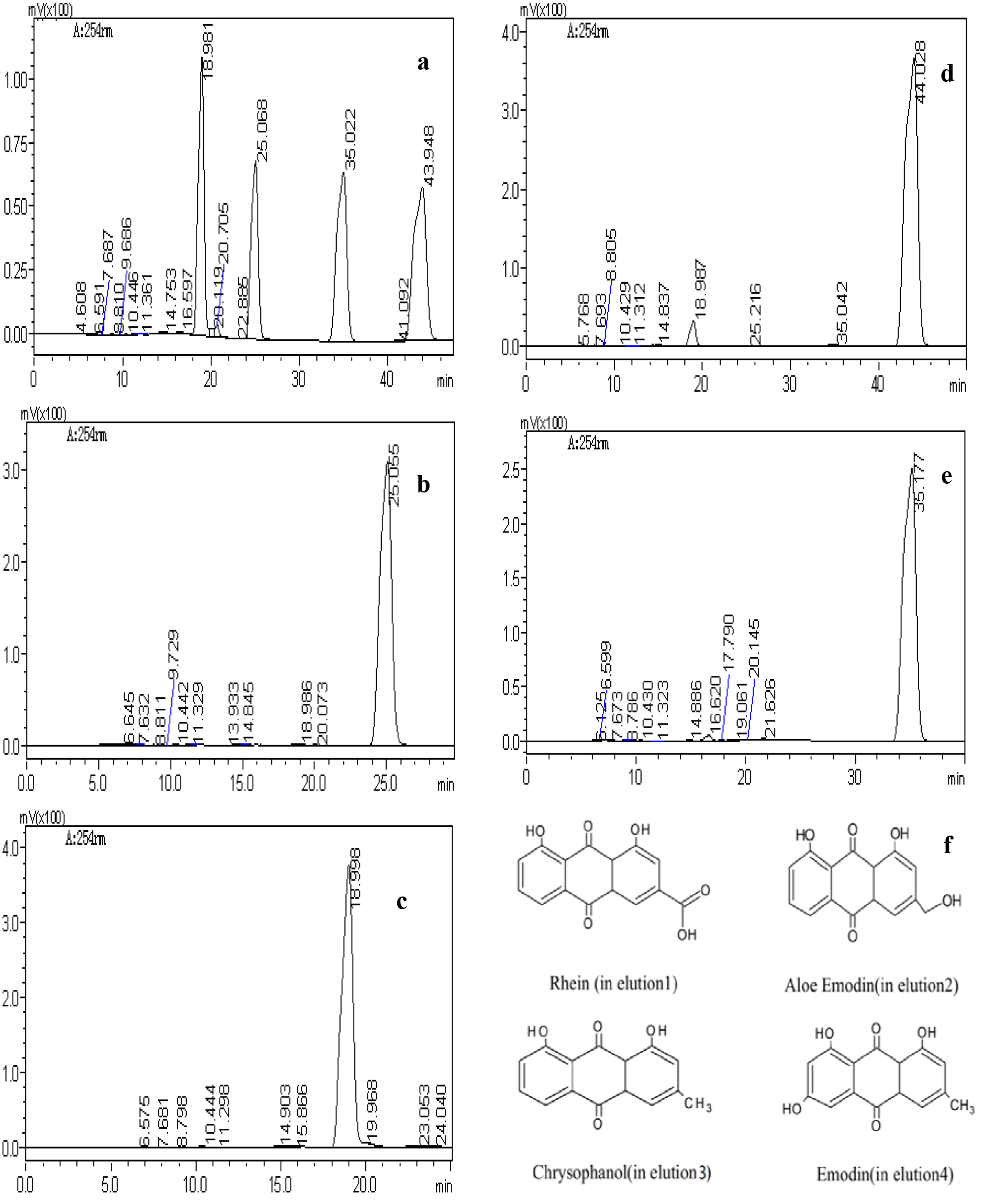
Liquid chromatogram for rust spots elutions; **a** Liquid chromatogram for standard sample; **b** Liquid chromatogram for rust spots elution 1; **c** Liquid chromatogram for rust spots elution 2; **d** Liquid chromatogram for rust spots elution 3; **e** Liquid chromatogram for rust spots elution 4; **f** Molecular structures of rhein, aloe-emodin, chrysophanol and emodin

It was all known that the chrysophanol belonged also to the anthraquinone compounds. The chrysophanol could fight availably against the apoptosis by means of improving the mitochondrial activity and the steady - state of the cell membrane, which was beneficial to the antiapoptotic impact of the hypoxia injury cells. After the hypoxia injury, there was a prominent variation for the permeability of the cell membrane on the cell surface. The augmentation of the cell damage degree promoted the reinforce of the permeability on the cell membrane, as well as the increase of the release quantity and synthetic level. A large number of the experimental results showed that the rhubarb phenol could improve the permeability of the cell membrane through the stability of the cell membrane, and stabilize the internal environment in the cell further, which reduced the damage level of the hypoxic cells significantly, enhanced the resistance ability on the hypoxia as for the cell self. In this experiment, the existence of the rhubarb phenol in the fragrant pear skin could improve significantly the permeability of the skin cell membrane, meanwhile stable the inner environment in the cell. Thus, the anoxic environment created by the controlled atmosphere storage could make the damage extent of the hypoxic cells on the fragrant pear skin reduce quickly to a minimum degree, which was favorable to enhance the cell’s ability to resist the hypoxia condition. Therefore, the rhubarb phenol had an advantage obviously role for the freshness of the korla fragrant pear, which could prolong the shelf life of the korla fragrant pear.

### LC - MS analysis of Elution 4

The fact, on the LC - MS analysis on Fig. 4 and the prototype HPLC information of Elution 4 in Fig. 5a and 5e, was agreed with the previous records for emodin (Ranjini *et al.*, 2011; Chen *et al.*, 2012)

As three elutions above, the emodin detected from the elution 4 was also a kind of the anthraquinone active substance with the typical structure of 9, 10 - anthraquinone. Therefore, the emodin displayed just more stablility at the highest oxidation levels. Previous studies had shown that the emodin had an important role in some aspects, such as the anti - cancer, anti - inflammatory, antibacterial, antiviral, anti - oxidation and purgative activity (Yan *et al.,* 2014). Antibacterial mechanism of the emodin had to do with the electron transfer inhibiting the mitochondrial respiratory chain and breath, including the oxidation and dehydrogenation of the metabolism intermediates from the amino acid, sugar, and protein. Emodin could inhibit the growth of the bacteria by adjusting the ultima synthesis of the nucleic acid and protein. In this experiment, the emodin on the korla fragrant pear skin could inhibit the electron transfer of the mitochondrial respiratory chain and the breath of the epidermal cells, including the oxidation and dehydrogenation of the metabolism intermediates from the amino acid, sugar, and protein of the pidermal cells, restrain the microbial activity of the korla fragrant pear skin eventually in order to achieve the goal of the preservation quality on fragrant pear.

In addition, emodin, as a natural inhibitors, had also certain inhibitory effect on the non - enzymatic glycosylation system. In the initial stage, the emodin could inhibit the condensation and cyclization of the carbonyl of the glucose and the amino of lysine from the fructosamine. The fructosamine after the amadori rearrangement was cracked into the dicarbonyl compounds promoting the obvious inhibitory effect on the glyoxal again after the enolization, which was a sign entering the middle stage of the non - enzymatic glycosylation system. Finally, the dicarbonyl compound and amino compound were polymerized into the final product by the polymerization of the aldehyde group and amino group. While the emodin had better inhibition role for these products including the carboxymethyl lysine (CML) and the melanoidin. In the process of the reaction, glyoxal, as a precursor of the carboxymethyl lysine, could be generated easily by the autoxidation of the glucose, or by the oxidation of the intermediate coming from the maillard reaction. So the emodin could suppress significantly the glycosylation through the antioxidation function. In addition, the emodin had no obvious inhibition influence to the intermediate, 5 - hydroxymethyl furfural, of the maillard reaction, which was confirmed further to the fact above. Thus, the emodin detected in this experiment endowed a strong antioxidant capacity to the skin of the korla fragrant pear, which could improve a capacity resisting the alternation of the external environment. In consequence, the existence of the emodin was conducive to the preservation of the fragrant pear.

In sum, the main bioactive constituents originating from the rust - colored substance collected from the pericarp of the Kurle pear were anthraquinones, namely aloe - emodin, rhein, emodin and chrysophanol (Fig. 5f), respectively, based on the LC - MS analysis combining with the studied results previously.

### Function of related anthraquinone

These anthraquinones synthesized via the polyketide pathway, such as emodin, chrysophanol, rhein, aloe - emodin, and physicione, were of the hydrophobic and a large number of the biological function, such as the anticancer (Shoemaker *et al.,* 2005; Huang *et al.,* 2007), antimicrobial and anti - inflammatory (Fosse *et al.,* 2004; Tseng *et al.,* 2006).

Research had demonstrated that the emodin was an effective inhibitor of the *neu* and *ras* oncogenes in vitro of kinase enzyme (Chang *et al.,* 1996), while emodin, together with anthraflavic acid, had also exhibited amicable protection mechanism against the benzo(a)pyrene mutagenicity and dietary pyrolysis mutagens through the Ames bacterial mutagenicity assay (Ayrton *et al.,* 1988; Lee *et al.,* 1991). It was interested that the orange aloe - emodin could induce the micronucleus frequencies in the mouse lymphoma L5178Y cells in the *in vitro* micronucleus test (Frisvad 1989), and the fungal chrysophanol were also considered as the mycotoxins owe to the violent genotoxicity today (Mueller *et al.,* 1998). Thus the detection of the emodin, chrysophanol, rhein, aloe - emodin and physicione suggested that some of these fungi possessed usually potent mycotoxigenic. Research result, however, about CaCl2 and pullulan treatments improving and even inhibiting the development degree of the brown spots on ‘Huangguan’ pear by delaying the loss of the polyphenol substances and maintaining the structural integrity of the cell membrane (Kou *et al.,* 2015) provided really a train of thoughts for the further research about the relationships between the pullulan and the aloe - emodin, rhein, emodin and chrysophanol. In this study, HPLC and LC - MS were used to analyze the rust spots on the pericarp of Kurle pears. As a result, four elutions (elutions 1 to 4), the material sources of the rust spots, were identified as rhein, aloe - emodin, chrysophanol and emodin, respectively, which was similiar to previous works(Kou *et al.,* 2015), and also provided a foundation for the future study of the rust spots.

### Ultrastructure of rusty spot and intact parts

The rusty spot parts of the korla fragrant pears had irregular black shadows, while the intact ones of which was white and transparent under the transmission electron microscopy (TEM) (Fig. 6). From Figs. 6d and 6e, the cell membrane and cell wall of the rusty spot sections was disappearing gradually with the serious damage of the cell wall, which led to the unclear cell contour ultimately. Therefore, there was not nearly a complete cell in the pears surface with the rusty spot. Another, the irregular black substance could be simultaneously and clearly found in the cell. Moreover, the ultrastructure of the avocado fruit cell had been observed during the mature process with the electron microscope. During the sclereid period, the intercellular layer, including each side ranged the filaments closely, of the fruit cells was clearly visible. However, the fruits accelerating the decrepitude promoted the gradual disintegration of the filaments and intercellular layer in the cell after the respiration peak (Ruth *et al.,* 1979; Chatelet *et al.,* 2008), which was similar to the some conclusions, such as the damage of the cellular structure, the autolysis of the intercellular layer, as well as the rusty spot being gradually formed on the korla fragrant pear after the respiratory climacteric during the postharvest storage. However, the important information supplied stemming from Fig. 6c and 6f showed that the cells were enough intact, oval and rowed neatly, and the outline of the cell wall was also distinct enough.

**Fig. 6.**
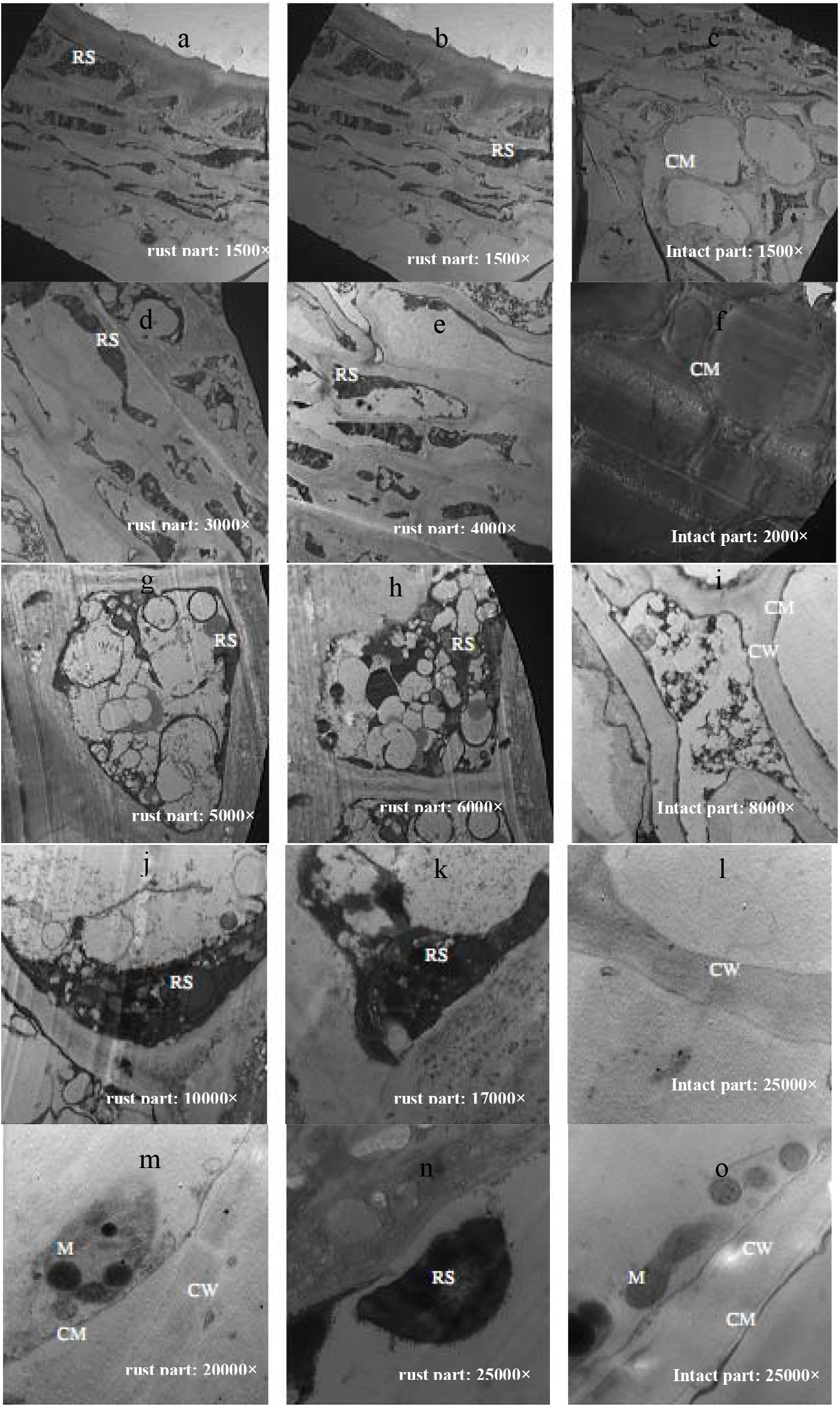
The cell ultrastructure of the rust and intact parts Note: RS-rust; CW-cell wall; CM-cell membrane; M-mitochondria

The black shadow, differ from that of the rusty spot section but integrated cell membrane and cell wall, of Fig. 6o was regular, which was induced by the mitochondria. The chrysophanol could adjust the mitochondria function, while the antibacterial mechanism of the emodin had to do with the electron transfer inhibiting the mitochondrial respiratory chain and breath, including the oxidation and dehydrogenation of the metabolism intermediates from the amino acid, sugar, and protein. However, the mitochondria of hwangkumbae, the clear double structure in internal envelope, and most of the round or oval shape, grew in the cell wall or next to a plasmid during the young fruit period (Cuthbertson *et al.,* 2003). From Fig. 6m, there were still part of the cell inclusions in the rust stain section of the korla fragrant pear, mitochondria, but without the organelles, such as the endoplasmic reticulum, golgi apparatus and microbody because of the mitochondrial more resistance collapse, could sometimes continue until the senescence phase, than the other micro. Meanwhile, the change of the mitochondrial membrane, such as the swelling, formed the cavity and destruction of the structure, led to gradually breaking apart of the mitochondria. Mitochondria were an important energy converter of the fruit cell. As we all known, the energy of the cells were supplied mainly by the mitochondria on the storage duration after the harvest. So, with the apolexis of the fruit, the fewer the number of the mitochondria was, the less the energy in cells would be, which abated correspondingly the repair and synthetic capacity of the cell, accelerated the collapse of the cell membrane, and even cell death finally (Chatelet *et al.,* 2008).

As could be seen from Fig. 6, there were black substances deposited in the skin cell during the formation of the rust stain. With the continued expansion of the pear rust stain area, these black materials began to extend gradually from the gap between the cells to the intracellular space, and even until the whole cell. At this time, the dark brown spots or patches in the pear skin were continuously extended likewise. It was interesting that during the late period of the pear storage, the number of the ribosome in rusty spot cells began to reduce gradually with the degeneration of the chloroplast badly destroyed, while the endoplasmic reticulum and golgi apparatus was rapidly disintegrated or collapsed owe to the vesiculation. At last, the tendency that the mitochondria began to collapse with the collapsed tonoplast before the microorgans were disintegrated completely became irreversible really, which led inevitably to the damage of the cytoplasm membrane further, and the death of the cells ultimately. Once the structure of the cell was destructed, some requirements, such as the dissociation of the intracellular phenols, as well as the increase of the phenylalanine ammonia enzyme and polyphenol oxidase activity, generated the dark brown substances under the interaction between these enzymes and phenolic substrates. With the destruction of the cell structure, these substance were still persistently increasing until the whole cell, which was the whole evolutionary process of the rusty spot development. The study of Cuthbertson and Murchie (Cuthbertson *et al.,* 2003) found that during the formation of the rusty spot on the hwangkumbae, the cell suberized stage by stage with the deepened color of the rusty spot gradually, but there was lack cause definitely now. Whereas we could see from the trial, there was a reasonable reason originating from the enzymatic browning. All in all, the combined action, namely both the stimulation under the adverse environment accelerated the maturation and aging of the pear, and the damage and collapse of the intracellular organelles to a variable extent, induced or launched the complicated physiological and biochemical process during the whole period.

The microstructure characteristic of Korla fragrant pear with the rusty spot and without the intact parts showed advantageously that the cytomembrane and cytoderm of the rusty spot location was bit by bit disappearing, while other organelles, such as the mitochondria, endoplasmic reticulum and golgi apparatus, were gradually disintegrated. As a result, the extended range and degree of the black deposits were increased abidingly from the local of the cells to the whole ones, it was without doubt which was an overt sign against the emergency response outside the environmental damage as for the skin of the fragrant pear, supported also by some conclusions on the eluent in the trial above (Fig. 1 – 5). But the fact we couldnot ignore was that the intact position was not an object of the microbial invasion, or not easy to be invaded by the microbes owe to the integral structures including some properties, such as the organizational hardness, structural material, enzyme activity of the peel, as well as some factors including the lower conditions of the environmental temperature created by the controlled atmosphere storage and the qualified sanitary condition. Therefore, there were no loss of the substances between and in cells, and the rust spots with the defensive function were not come into being on the surface of the peel (Fig. 6c, f, i, l, and o). Naturally, There were not four anthraquinones, including aloe - emodin, rhein, emodin and chrysophanol on the surface of the unbroken peel. Thus, not only the damage of the cell structure on the pear skin, but also the increase of the lignin and secondary protection organization of the intercellular layer, including the further lignification of the secondary organization led to the formation of the rusty spot finally, which postponed the senescence of the pear fruit further.

## Conclusion

In conclusion, the main rust - colored substance originating from the bioactive constituents collected from the pericarp of Kurle pear were anthraquinones, including aloe - emodin, rhein, emodin and chrysophanol. Another, the microstructure information obtained from the rusty spot and intact parts in Korla fragrant pear drew a prominent conclusion that the gradual variation of some organelles, such as the cytomembrane, cytoderm, mitochondria, and endoplasmic reticulum of the rusty spot location resulted in the abiding expansion of more and more black deposits toward the whole cells. The damage of the cell structure on the pear skin, as well as the increase of the lignin and secondary protection organization, including the further lignification organization, in the intercellular layer, led to the formation of the rusty spot finally, which was the result of the self - defence against the harsh environment.

